# Exacerbated salmonellosis in poly(ADP-ribose) polymerase 14 deficient mice

**DOI:** 10.1101/2025.01.01.630990

**Authors:** Madhukar Vedantham, Lauri Polari, Tiia Rissanen, Arto Tapio Pulliainen

## Abstract

*Salmonella enterica* subspecies *enterica* serovar Typhimurium (*S.* Typhimurium) is an enteropathogen annually causing millions of acute infections ranging from gastroenteritis to life-threatening invasive systemic disease. Strong mucosal inflammation is characteristic for the infection driven by the innate immune receptor activation but also directly by the invading bacterium. It remains unknown how and at which point the mucosal inflammation turns anti-bacterial and how the tissue homeostasis is restored. Here, we investigated the expression and function of poly(ADP-ribose)polymerase (Parp14), a known cytoplasmic and nuclear regulatory protein of immune cells, in the mouse streptomycin-pretreatment model of *S.* Typhimurium infection. In the infected mice, Parp14 expressing cells, some of which were macrophages, were detected throughout the gastrointestinal tract. However, the most strongly Parp14 expressing cells were the epithelial cells. Based on small intestine single cell RNA-Seq analysis of different epithelial cell types, the expression of *parp14* was pronounced in the enterocytes and Tuft cells. Mice with a body-wide genetic deficiency of Parp14 suffered from exacerbated *S.* Typhimurium colitis. The histopathological analysis of the large intestine revealed increased immune cell infiltration, Goblet cell loss, and epithelial erosion. At the same time lower numbers of viable bacteria were detected. Based on a bulk tissue RNA-Seq analysis, transcriptional signatures either missing or down-regulated in the infected Parp14 deficient mice were detected. These signatures were enriched with genes related to cell adhesion, cell division and cytoskeletal rearrangements, and genes related to infection and immune responses. It appears that Parp14 has potential functions in the regulation of tissue architecture and mucosal inflammation in the large intestine of *S.* Typhimurium infected mice.

## INTRODUCTION

*Salmonella enterica* subspecies *enterica* serovar Typhimurium (hereafter *S.* Typhimurium) is a food- and waterborne enteropathogen annually causing millions of acute infections ranging from the self-limiting non-invasive gastroenteritis to life-threatening invasive systemic disease (1, 2). The globally emerging antibiotic-resistant strains complicate the clinical management of the most severe forms of *S.* Typhimurium infection (2–4). Based on the analyses of patient tissue samples, *S.* Typhimurium elicits mucosal inflammation, in particular, in the terminal ileum and colon characterized by a massive neutrophil influx (5, 6). It is believed that the expression of bacterial virulence factors (7, 8), such as flagella and type I fimbriae, drive the initial contacts between *S.* Typhimurium and colon epithelial cells, the enterocytes, through the protective barrier created by the colon lumen microbiota (9) and dense epithelial cell mucous layer (10). It appears evident that *S.* Typhimurium also actively interact with trans-epithelial and/or lamina propria-associated dendritic cells as well as macrophages (7, 8). Upon initial contacts with the host cells, *S.* Typhimurium is thought to utilize two different type 3 secretion systems (TTSS-1 and TTSS-2) and a plethora of TTSS effector proteins, to drive deeper colon wall invasion and intracellular as well as extracellular replication (7, 8). Some of the TTSS effects are counterintuitively driving mucosal inflammation, which is additionally activated by the classical innate immunity receptors such as Toll-like receptors (TLRs) recognizing pathogen associated molecular patterns (7, 8, 11, 12). It is believed that *S.* Typhimurium benefits from the colon inflammation to outcompete the colon lumen microbiota (13, 14), that is, via creation of gut dysbiosis, and to obtain nutrients boosting the colon lumen bacterial replication (15). It remains unknown how and at which point the mucosal inflammation turns anti-bacterial and how the tissue homeostasis is restored. Elucidation of molecular mechanisms regulating the mucosal inflammation in *S.* Typhimurium infection could be useful for the development of novel host-targeted anti-bacterial pharmaceuticals.

Eukaryotic cells rely on dynamic cell signaling events to mount responses to external perturbations, such as an invading bacterial pathogen. Post-translational modifications influence the location, half-life, and activity of proteins, and thereby serve as a powerful regulatory mechanism of cell signaling. Poly (ADP-ribose) polymerase 14 (Parp14) is a multidomain ADP-ribosyltransferase (ART) enzyme of the Parp protein family (16). Parp14 was identified as a Stat6 interacting protein (17) with a postulated role as a transcriptional co-factor in interleukin-4 (IL-4)-induced Stat6-dependent gene expression (17–20). Parp14 catalyzes protein ADP-ribosylation, including itself, (21–23), which refers to the covalent conjugation of an ADP-ribose moiety from nicotinamide adenine dinucleotide (NAD+) onto the substrate protein with simultaneous release of nicotinamide (16). Parp14 functions that are independent of the ART activity such as scaffolding and modification reversal also appear possible, as exemplified by the recent work on ADP-ribose binding and ADP-ribose hydrolysis activities of the Parp14 macrodomains (24, 25). The functional outcomes of Parp14-catalyzed protein ADP-ribosylation have remained largely unknown. Regulation of macrophage activation via ADP-ribosylation of Stat-proteins has been proposed (26, 27). The recent work on identification of Parp14 as the key ART of the interferon-γ (IFN-γ)-inducible protein ADP-ribosylation response is expected to pave the way for a better understanding of the cellular and physiological functions of Parp14 (22, 28–31). It is plausible that Parp14 has functional relevance to mount a controlled physiological response to a bacterial infection. However, to the best of our knowledge, only one *in vitro* phenotypic study employing Parp14 depletion has been published (32). The Parp14 deficient RAW264.7 macrophages contained more viable *S*. Typhimurium bacteria, and, upon *S*. Typhimurium infection, produced less microbicidal nitric oxide as well as had a defective expression pattern of a number of inflammation-related genes, such as *ifnb1*, *ccl5*, *cxcl10*, and *ifit1* (32). Therefore, Parp14 appears as a protein with functional importance in the anti-bacterial responses of macrophages.

We hypothesized that Parp14 is involved in the anti-bacterial mucosal immune response, and, if so, its malfunction could play a role in the development of bacterial gastroenteritis. We explored the expression of Parp14 and the effect of its genetic deficiency in the mouse streptomycin-pretreatment model of *S.* Typhimurium infection (8, 33). Parp14 was expressed by macrophages but, in particular, by the epithelial cells throughout the mouse gastrointestinal tract. An exacerbated salmonellosis was witnessed in the Parp14 deficient mice. This phenotype paralleled with missing or down-regulated transcriptional signatures with functional relevance in the regulation of tissue architecture and mucosal inflammation.

## MATERIALS AND METHODS

### Mouse experimentation

National Animal Experiment Board has approved our mouse experiments in C57BL/6N background (ESAVI/24418/2018). **i) Colony breeding**. Our colony of Parp14 deficient mice was established based on the previously described body-wide Parp14 knockout mice (34), kindly provided by Adam Hurlstone (University of Manchester, UK), in a specific-pathogen-free area at Central Animal Laboratory of the University of Turku with free access to soy-free diet and water ad libitum. The Parp14 knockout mice were backcrossed for 10 generations to C57BL/6N background before starting the experiments (35). **ii) *Salmonella* infection.** Experiments with female C57BL/6N mice of age 6-8 weeks were performed as described (33). The mice were allocated to two groups, i.e., PBS control and *Salmonella* infection groups, with similar starting body weights. The naturally streptomycin-resistant SL1344 strain of *S.* Typhimurium was purchased from the Culture Collection University of Gothenburg (CCUG 51871), Gothenburg, Sweden. Bacteria were grown for 12h at 37°C in Luria-Bertani (LB) medium with shaking, diluted 1:20 in fresh medium, and sub-cultured for 4h with shaking. Bacteria were washed twice and suspended in ice-cold sterile phosphate buffered saline (PBS). Water and food were withdrawn 4h before per os (p.o.) treatment with 20 mg of streptomycin (75 μl of sterile solution or 75 μl of sterile water). Afterward, animals were supplied with water and food ad libitum. At 20 h after streptomycin treatment, water and food were withdrawn again for 4 h before the mice were orally cavaged with 10^8^ colony forming unit (CFU) of *Salmonella* (50 μl suspension in PBS p.o.) or with PBS (50 μl). Thereafter, drinking water ad libitum was offered immediately and food 2h post-infection (p.i.). At the indicated times p.i., mice were sacrificed by CO_2_ asphyxiation, organs were weighed, colon length was measured, and tissue samples were processed for bacterial viability quantitation (CFU/g) (samples collected in cold PBS), histological analyses (samples collected in 4% paraformaldehyde) and RNA analysis (samples collected in liquid nitrogen).

### Quantitation infection severity

**i) Enumeration of viable bacteria.** Fecal pellets were placed in 500 μl of ice-cold PBS and suspended to homogeneity on ice by vortexing and pipetting. The distal small intestine, cecum, proximal and mid large intestine, mesenteric lymph nodes, spleens, and livers were removed aseptically and homogenized in ice-cold PBS at +4°C by using stainless steel balls (IKA 5mm stainless steel balls, Fisher scientific) and compact bead mill (TissueLyser LT, Qiagen). The CFUs were determined by plating different dilutions on LB agar plates (streptomycin, 50 μg/ml). Plates were incubated at 37°C for approximately 12h before counting the colonies. **ii) Histopathologial analysis.** Formalin-fixed paraffin-embedded (FFPE) tissues were cut in longitudinal 5 µm thick sections prior to hematoxylin & eosin (HE) staining. HE staining was performed using standard methods. All samples were scanned using a Pannoramic 1000 Slide scanner (3DHistech, Budapest, Hungary) with 20x objective and analyzed with a Pannoramic viewer (3D Histech, software version 1.15.4). Histopathology was scored based on four variables, that is, i) immune cell infiltration to the gut wall as 0 (healthy/neglectable numbers of immune cells), 1 (some immune cells), 2 (moderate level of immune cells), 3 (high level of immune cells indicative of a severe inflammation); ii) edema of the gut wall as 0 (healthy/neglectable edema), 1 (some edema), 2 (moderate edema), 3 (strong edema indicative of a severe inflammation); iii) epithelial erosion at the gut wall as 0 (healthy/neglectable erosion), 1 (mild loss of epithelial cells), 2 (moderate loss of epithelial cells), 3 (strong loss of epithelial cells, and erosion going through muscular lamina indicative of a severe inflammation); iv) loss of Goblet cells at the gut wall as 0 (healthy/neglectable loss of Goblet cells), 1 (some loss of Goblet cells), 2 (moderate loss of Goblet cells), 3 (severe loss of Goblet cells indicative of a severe inflammation). Two people performed the scoring independently and blind for the animal groups to obtain the average scores for statistics. **iii) Statistical analyses.** The statistical analysis for studying association of variables (tissue variables, histology scores, bacterial load variables, weights) with mice group [wild-type (wt) vs. Parp14-KO] was analyzed separately by termination day. Tissue variables (spleen, liver, colon length) and bacterial load variables (proximal colon, liver, spleen, MLNs, fecal pellets, distal small intestine) were summarized with descriptive statistics and associations between mice group were studied with Kruskal-Wallis test. Histology scores (edema, immune infiltration, goblet cell loss, erosion) were measured by distal small intestine, proximal colon and whole cecum. Associations between mice group and variables were also studied by Kruskal-Wallis test. Weight percentages were summarized with descriptive statistics. Differences between groups in the 5-day follow-up were analyzed using the Friedman test because the normality assumption was not met. Wilcoxon signed rank test was used to study the difference of days. The normality of variables was evaluated visually and tested with the Shapiro-Wilk test. Due to the non-normality of the continuous variables, nonparametric methods were used. All tests were performed as two-tailed and the statistical significance level was set at 0.05. The analyses were carried out using the SAS system, version 9.4 for Windows (SAS Institute Inc., Cary, NC, USA).

### Parp14 immunohistochemistry

The 5 µm thick FFPE sections were stained for Parp14 with mouse monoclonal antibody (sc377150, Santa Cruz Biotechnology, dilution 1:500) and detected using the mouse specific HRP-DAB (ABC) detection immunohistochemistry (IHC) kit (ab64259, Abcam). Tissue sections were air dried for 2 h at room temperature, placed in 37°C incubator overnight, deparaffinized in xylene and rehydrated with alcohol gradients. Endogenous peroxidase was blocked with ‘Peroxidase blocking solution’ provided with the IHC kit. The sections were immersed in prewarmed 10 mM Na-citrate buffer (freshly prepared, pH 6.0) and kept in boiling water bath for 20 minutes. After antigen retrieval, sections were rinsed in PBST [phosphate buffered saline (PBS) with 0.01% Tween-20] followed by BSA blocking (5% w/v in PBST) for 1 h at room temperature to reduce nonspecific binding of the antibodies. The primary anti-Parp14 antibody in BSA (5% w/v in PBST) was added to tissue sections and incubated overnight at 4°C in a humidified chamber. Post incubation with primary antibody, tissue sections were rinsed in PBST twice for 5min each and incubated with biotinylated anti-mouse secondary antibody (provided with IHC kit) for 1h at room temperature. Streptavidin-HRP conjugate was added and then stained using 3-3’ Diaminobenzidine (DAB) as a chromogen (both solutions were provided with IHC kit). Harris hematoxylin was used as a nuclear counterstain. Sections without incubation of primary antibody served as negative controls. Mounting was done using Histo-Clear and sections were allowed to sit at room temperature for 12h before imaging. Imaging was done using Zeiss AxioImager M1 microscope with 5x, 20x, 40x oil or 63x oil objective lenses.

### Parp14 and F4/80 double immunofluorescence staining

Staining was performed as previously described (35). Briefly, Alexa-fluor 647 conjugated anti-F4/80 (MCA497A647, Biorad, dilution 1:500) and anti-Parp14 (sc377150, Santa Cruz Biotechnology, dilution 1:500) were mixed in 5% (w/v) BSA in PBST. Secondary antibody Alexa-fluor 488 goat anti-mouse IgG (H+L) (A11001, Invitrogen, dilution 1:1000) was used to visualize the anti-Parp14 antibodies. Mounting was done using Histo-Clear post DAPI counterstaining and sections were kept in dark at 4°C before Zeiss AxioImager M1 imaging.

### QuPath-based quantitation of Parp14 immunohistochemical staining

The anti-Parp14 stained tissue sections of wt mice (distal small intestine, cecum, and proximal large intestine) were scanned using a Pannoramic 1000 Slide scanner (3DHistech, Budapest, Hungary) with 40x objective and analyzed with a Pannoramic viewer (3D Histech, software version 1.15.4). Scanned slides were converted to .mrxs file type and specific project in QuPath (qupath.github.io, software version 0.5.1, (36)) was created to quantify the DAB OD (optical density). Parameters for stain vectors of Hematoxylin and DAB were set using automatic estimation in QuPath. For cellular detection, nucleus parameters used were: background radius - 8µm, minimum area −10µm^2^ and maximum area - 400µm^2^. Cell expansion was set at 5µm. From each tissue of each section, minimum 50 to maximum 200 horizontal villus crypts were selected for cellular detection and quantifying DAB staining intensity. Detected annotations were exported and statistical analyses were conducted using the one-way ANOVA with Tukey’s multiple comparison test to compare the means.

### Single cell RNA-Seq

The already published single cell RNA-Seq data on epithelial cell-enriched cell suspensions of control (4 mice) vs. *S. Typhimurium* infected (SL1344 strain, 2 days post-infection, 2 mice) C57BL/6J mice (37), was re-analyzed. We used the single cell RNA-Seq data analysis and visualization interface at the Broad Institute Single Cell Portal (https://singlecell.broadinstitute.org/single_cell).

### Isolation of RNA and quantitative PCR analysis

Total RNA was isolated from mouse tissues using TRIsure reagent (BIO-38033, Bioline Gmbh, Germany) and genomic DNA was digested using RNase-free DNase (rDNase) from Machery-Nagel as per the manufacturer’s instructions. Briefly, colon tissue was homogenized using stainless steel balls (IKA 5mm stainless steel balls, Fisher scientific) and compact bead mill (TissueLyser LT, Qiagen). Colon tissue was placed in TRIsure reagent during homogenization. After this, chloroform was used for phase separation. RNA in the upper aqueous phase was precipitated using isopropyl alcohol. Pellet was washed with 70% ethanol and dissolved in nuclease free water. Dissolved RNA was mixed with rDNase and rDNase reaction buffer as per manufacturer’s instructions. This mixture was incubated at 37°C for 10 min. Post gDNA digestion, RNA was precipitated using 3M sodium acetate, pH 5.2 and 96% ethanol. Pellet was washed with 70% ethanol and dissolved in RNase free water. RNA purity and concentration were measured using DeNovix DS-11 spectrophotometer (Wilmington, DE, USA). The dissolved RNA (1µg) was reverse transcribed using SuperScript III reverse transcriptase (#1808044, ThermoFisher Scientific) and Oligo (dT) 12-18 Primer (#18418012, ThermoFisher Scientific). Separate real-time PCR (RT-PCR) was carried out in duplicates using TaqMan gene expression assays (Applied Bio systems) for *parp14* (Assay ID: Mm00520984_m1), *il1b* (Assay ID: Mm00434228_m1), *il6* (Assay ID: Mm00446190_m1), *ccl2* (Assay ID: Mm00441242_m1), *tnfa* (Assay ID: Mm00443258_m1) and the reference gene, glyceraldehyde-3-phosphate dehydrogenase, *gapdh* (Assay ID: Mm99999915_g1) on the Rotor-Gene Q real-time PCR cycler (Qiagen). Thermal cycling conditions included an initial denaturation step at 95°C for 10 min followed by 40 cycles of 95°C for 15s and 60°C for 1 min. Relative mRNA levels were determined using the 2^-ΔΔCT^ method with GAPDH as reference (38). If the standard deviation of duplicate Ct values from a sample was 0.5 or more, results of those samples were not used for statistical analysis. Statistical analysis of 2^-ΔΔCT^ values was done using the unpaired t test (two tailed).

### RNA-Seq runs and data analysis

Total RNA of the distal colon tissue samples was extracted as described above. **i) RNA quality.** RNA integrity, including all the other RNA-Seq wet-lab techniques described below, was assessed by Novogene Co., LTD (Cambridge, UK) using the RNA Nano 6000 Assay Kit of the Bioanalyzer 2100 system (Agilent Technologies, CA, USA). **ii) Library preparation for transcriptome sequencing.** Briefly, mRNA was purified from total RNA using poly-T oligo-attached magnetic beads. Fragmentation was carried out using divalent cations under elevated temperature in First Strand Synthesis Reaction Buffer (5X). First strand cDNA was synthesized using random hexamer primer and M-MuLV Reverse Transcriptase (RNase H-). Second strand cDNA synthesis was subsequently performed using DNA Polymerase I and RNase H. Remaining overhangs were converted into blunt ends via exonuclease/polymerase activities. After adenylation of 3’ ends of DNA fragments, adaptors with hairpin loop structure were ligated to prepare for hybridization. To select cDNA fragments of preferentially 370-420 bp in length, the library fragments were purified with AMPure XP system (Beckman Coulter, Beverly, USA). Then PCR was performed with Phusion High-Fidelity DNA polymerase, Universal PCR primers and Index (X) Primer. At last, PCR products were purified (AMPure XP system) and library quality was assessed on the Agilent Bioanalyzer 2100 system. **iii) Clustering and sequencing.** The clustering of the index-coded samples was performed on a cBot Cluster Generation System using TruSeq PE Cluster Kit v3-cBot-HS (Illumia) according to the manufacturer’s instructions. After cluster generation, the library preparations were sequenced on an Illumina Novaseq platform and 150 bp paired-end reads were generated. The raw RNA-Seq data has been deposited to NCBI (https://www.ncbi.nlm.nih.gov) Gene Expression Omnibus (GEO) database with accession number GSE284287. **iv) Data analysis - Quality control.** Raw data (raw reads) of fastq format were firstly processed through Novogene’s in-house perl scripts. In this step, clean data (clean reads) were obtained by removing reads containing adapter, reads containing ploy-N and low quality reads from raw data. At the same time, Q20, Q30 and GC content the clean data were calculated. All the downstream analyses were based on the clean data with high quality. **v) Data analysis - Reads mapping to the reference genome.** Reference genome and gene model annotation files were downloaded from genome website directly. Index of the reference genome was built using Hisat2 v2.0.5 and paired-end clean reads were aligned to the reference genome using Hisat2 v2.0.5. We selected Hisat2 as the mapping tool for that Hisat2 can generate a database of splice junctions based on the gene model annotation file and thus a better mapping result than other non-splice mapping tools. **vi) Data analysis - Quantification of gene expression level.** The featureCounts v1.5.0-p3 was used to count the reads numbers mapped to each gene. Subsequently, Fragments Per Kilobase of transcript sequence per Millions base pairs sequenced (FPKM) of each gene was calculated based on the length of the gene and reads count mapped to this gene. **vii) Data analysis - Differential expression analysis.** Differential expression analysis of two conditions/groups (two biological replicates per condition) was performed using the DESeq2 R package (1.20.0). DESeq2 provide statistical routines for determining differential expression in digital gene expression data using a model based on the negative binomial distribution. The resulting *p*-values were adjusted using the Benjamini and Hochberg’s approach for controlling the false discovery rate. Genes with an adjusted *p*-value <0.05 found by DESeq2 were assigned as differentially expressed. **viii) Data analysis - GO and KEGG analyses.** To analyze cellular and physiological associations of the uniquely expressed genes (FPKM>1) and the differentially expressed genes (DEGs), we performed the Gene Ontology (GO) enrichment at the GO consortium website (https://geneontology.org) (39–41). In parallel, we performed the Kyoto Encyclopedia of Genes and Genomes (KEGG) pathway analysis (42–44) with SRplot (45) (http://www.bioinformatics.com.cn/srplot).

## RESULTS

### Parp14 is expressed by macrophages and epithelial cells in the mouse gastrointestinal tract

Littermates of female wt and Parp14 deficient mice (34, 35) were either orally gavaged with PBS or with *S.* Typhimurium strain SL1344 followed by sampling as described in Fig. 1A-B. First, we analyzed the expression of Parp14 at the protein level in wt mouse using a commercial anti-Parp14 monoclonal antibody, validated for epitope specificity in our previous study (35). We detected cells in the PBS and *S.* Typhimurium gavaged mice that were positive for Parp14 in the lamina propria of small intestine, cecum and large intestine (Fig. S1). Based on the double immunofluorescence staining with a monoclonal antibody recognizing the classical F4/80 macrophage marker, some of these Parp14 positive cells were macrophages as exemplified with the large intestine of infected mice (Fig. S2). In fact, it appeared that the most intensely Parp14-staining F4/80 positive cells were frequently embedded in the epithelial cell layer with apparent protrusions extending towards the gut lumen (Fig. S2). The most evident expression of Parp14, however, was witnessed in the epithelial cells in the small intestine, cecum and large intestine (Fig. S1). We executed a QuPath-based quantitation of the Parp14 staining intensity with entire tissue sections by selecting 50-200 horizontal villus cross-sections and thereby thousands of individual epithelial cells per animal (Fig. S3). This analysis revealed a temporal pattern of Parp14 staining (Fig. 2A). The Parp14 staining in the small intestine increased by the infection at day 1 but decreased lower than the resting level at day 5. The Parp14 staining in the cecum decreased by the infection at day 1 and from that level even further down at day 5. The large intestine showed a more complex staining pattern. The Parp14 staining increased by the infection at day 1 and from that level even further up at day 5 when the whole cell or the cytosolic QuPath readouts were compared. However, less Parp14 staining was detected in the nucleus at day 5 as compared to day 1. Next, we conducted a qPCR-based quantitation of *parp14* expression in all the three tissues. We did not detect statistically significant differences between the two time points (Fig. 2B). To substantiate our bulk tissue findings, we analyzed the already published single cell RNA-Seq data on epithelial cell-enriched cell suspensions of control vs. *S.* Typhimurium (SL1344 strain, 2 days post-infection) C57BL/6J mice (37). Out of the 8 detected small intestine epithelial cell sub-types (endocrine cells, enterocytes, enterocyte progenitors, Goblet cells, stem cells, transit amplifying cells, early transit amplifying cells, Tuft cells), the expression of *parp14* was pronounced in the enterocytes and Tuft cells of the infected mice (Fig. S4). Such a pattern of expression was not detected in the PBS gavaged mice. Of note, the expression pattern of the founding member of the Parp-family, *parp1* (16), was similar in the 8 epithelial cell types of PBS and *S.* Typhimurium gavaged mice being most pronounced in the endocrine cells. Taken together, Parp14 was expressed by the macrophages, but in particular, by the epithelial cells across the mouse gastrointestinal tract with a temporal pattern in *S.* Typhimurium infection.

**Figure 1.**
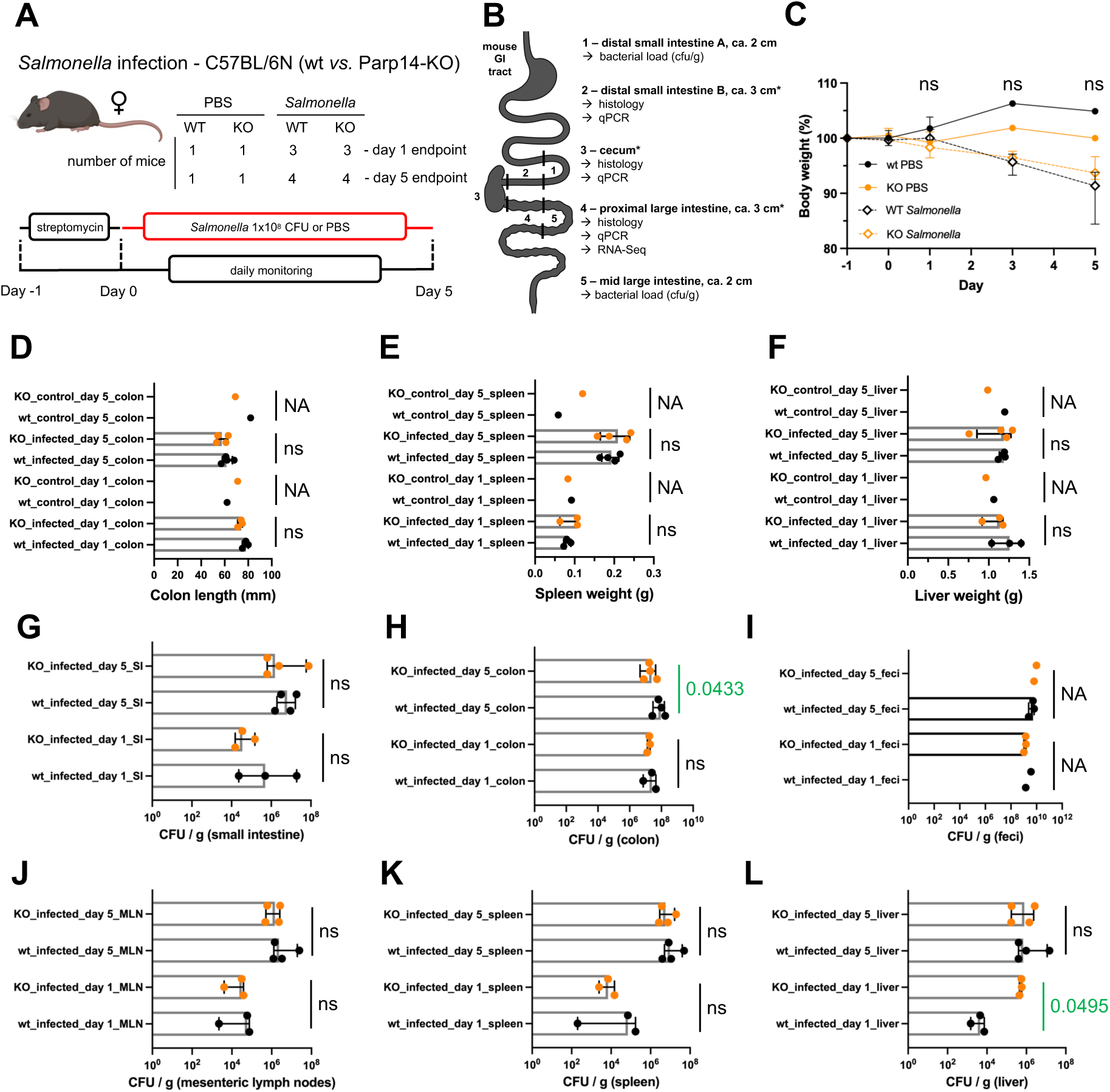
Minor macroscopic effects of Parp14 deficiency on the severity of salmonellosis. **A-B)** Schematic representation of the animal experimentation, sampling and executed analyses. The tissues marked with an asterisk were longitudinally cut in two pieces, one for histology and one for qPCR/RNA-Seq. Images were partially created with BioRender.com. **C)** Weight change of the mice during the course of the experiment relative to day −1 (medians with interquartile range). No statistical differences between the infected wt and Parp14 deficient mice were detected. **D)** Colon lengths at day 1 and day 5. **E)** Spleen weights at day 1 and day 5. **F)** Liver weights at day 1 and day 5. **G-L)** Determination of viable bacteria in different tissues at day 1 and day 5. The bars in sub-panels D-L are represented as medians with interquartile range. All individual data points are shown. Fecal pellets were not obtained from all of the mice. Statistical significance values for the differences between the infected wt and Parp14 deficient mice are shown in each D-L sub-panel (NA – not applicable, less than 3 datapoints/animal to compare; ns – not significant; significant with a green font, *p*<0.05).

**Figure 2.**
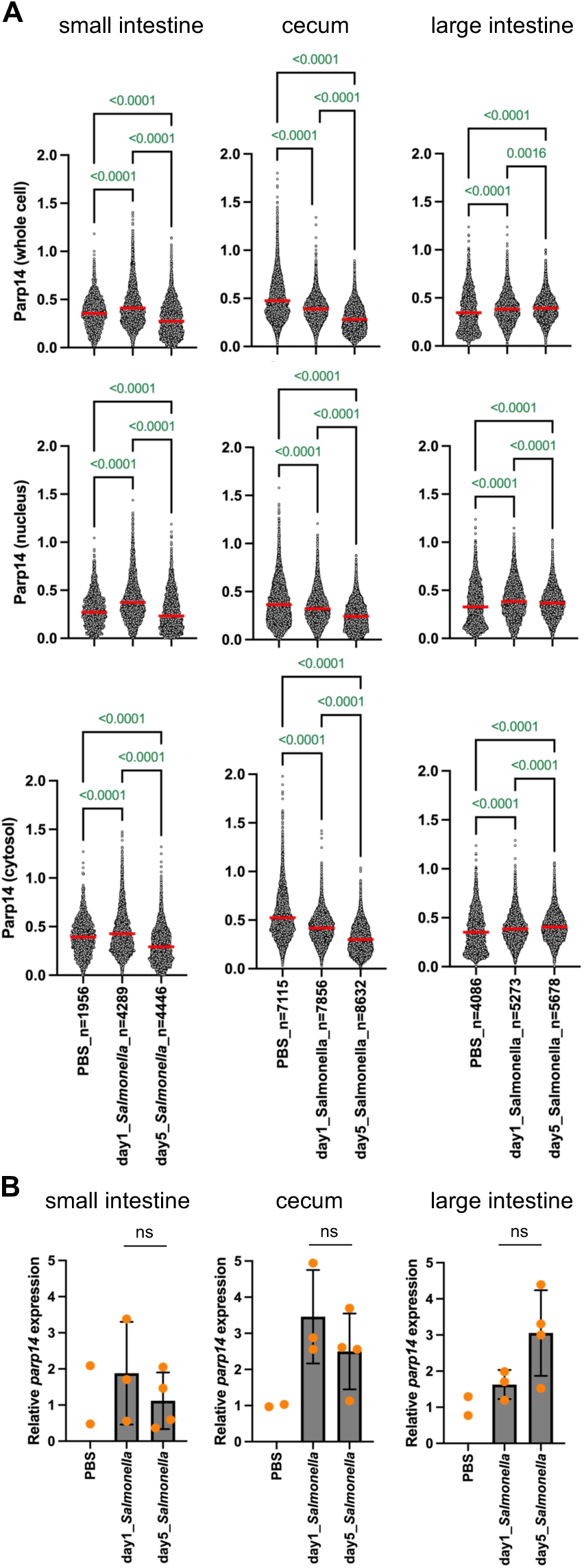
Quantitation of Parp14 expression in the mouse gastrointestinal tract. **A)** QuPath-based quantitation of Parp14 expression in the FFPE tissue sections. Examples of the Parp14 staining are shown in Fig. S1. One *Salmonella* infected day 5 mouse was left out from the quantitation due to poor quality of the FFPE tissue block. The values on the Y-axis refer to the means of DAB staining intensity, that is, DAB OD mean in the QuPath data output. Each dot refers to a single cell. The numbers of analyzed cells are indicated in the Y-axis. The red lines on top of the data points refer to the mean values. Statistical analyses were conducted using the one-way ANOVA with Tukey’s multiple comparison test to compare the means. **B)** The qPCR data on relative *parp14* expression (means with standard deviation, statistics with two-tailed unpaired t-test). Samples were included in the data analysis if they passed the 0.5 standard deviation Ct filter for replicate runs. No statistical analyses were executed against the PBS groups because there were less than 3 datapoints/animal to compare. The calibrators in each sub-panels are the mean dCq-values of the PBS samples.

### Minor macroscopic effects of Parp14 deficiency on the severity of salmonellosis

We monitored the PBS and *S.* Typhimurium gavaged mice daily also by quantifying their body weight up to the day 5 termination point. Infection caused weight loss in wt and Parp14 deficient mice in a statistically similar manner (Fig. 1C). No differences between the infected wt and Parp14 deficient mice were detected in the colon length, spleen weight or liver weight at day 1 or day 5 (Fig. 1D-F). However, when we quantified the numbers of viable bacteria in different tissues, some statistically significant differences were detected (Fig. 1G-L). The bacterial load (CFU/g) was higher in the liver of Parp14 deficient mice at day 1. In contrast, the bacterial load was lower in the colon of Parp14 deficient mice at day 5. Otherwise, no differences in the bacterial loads were detected. It appears, based on the measured macroscopic variables, that the effect of Parp14 deficiency on the severity of salmonellosis is minor.

### Exacerbated gastrointestinal histopathology in *S*. Typhimurium infected Parp14 deficient mice

We quantified epithelial erosion, tissue edema, tissue immune cell infiltration and Goblet cell loss in small intestine, cecum and large intestine from the H&E-stained FFPE sections (Fig. 3). Of note, the absence of Parp14 did not cause apparent small intestine, cecum or large intestine deformations in the resting state, as evidenced with the PBS gavaged mice (Fig. 2). This is in line with our previous comparative results with wt and Parp14 deficient male (35). Several statistically significant histological differences between the infected wt and Parp14 deficient were detected (Fig. 3). At day 1, the epithelial erosion was stronger in the small intestine of Parp14 deficient mice. At day 5, the epithelial erosion was also stronger in the large intestine of Parp14 deficient mice. Moreover, at day 5, the immune cell infiltration and Goblet cell loss were stronger in the large intestine of Parp14 deficient mice. It appears that the deficiency of Parp14 had sensitized mice to salmonellosis, as evidenced with the exacerbated histopathology, in particular, in the large intestine.

**Figure 3.**
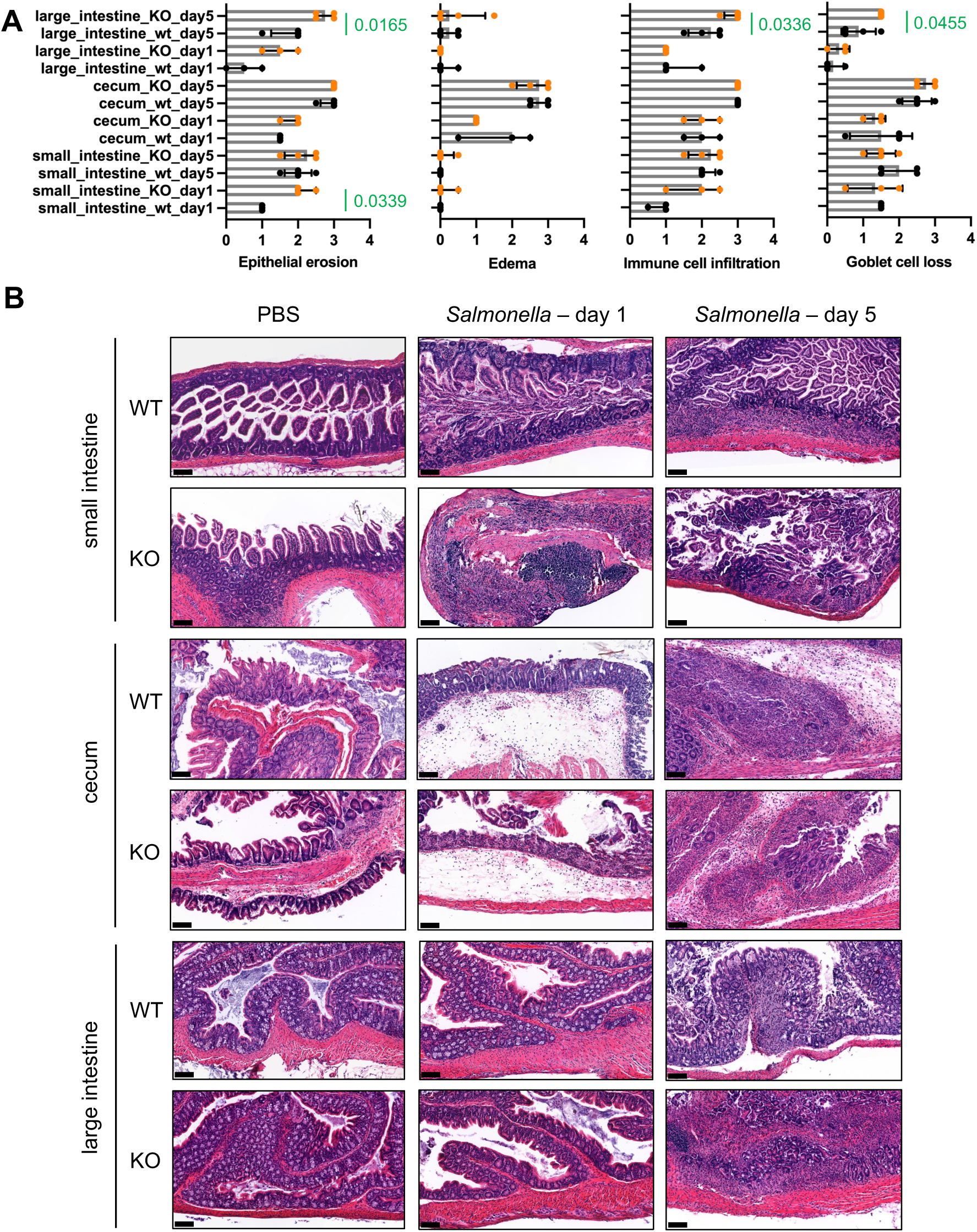
Exacerbated gastrointestinal histopathology in *S*. Typhimurium infected Parp14 deficient mice. **A)** Quantitation of histopathological variables, that is, epithelial erosion, tissue edema, immune cell infiltration and Goblet cell loss in distal small intestine, cecum, and large intestine (medians with interquartile range). Statistical significance for the differences between the infected wt and Parp14 deficient mice are shown in each sub-panel (significant with a green font, *p*<0.05). **B)** Representative H&E-stained tissue sections. 15x air objective images are shown. The size of the scale bar in each sub-panel refers to 100 µm.

### Transcriptional signatures uniquely detected in large intestines of *Salmonella*-infected wt and Parp14 deficient mice

We performed a bulk tissue large intestine RNA-Seq analysis using triplicate samples of day 1 infected wt and Parp14 deficient mice. First, we looked on the identities of genes detected to be expressed using the canonical transcript detection cut-off FPKM (Fragments Per Kilobase per Million mapped fragments)-value of >1. As shown in Fig. 4A and Suppl. data file 1_1-3, 11648 genes were detected in both genotypes, as well as 520 and 325 genes specifically in the wt and Parp14 deficient mice, respectively. Next, we ran Gene Ontology (GO) term searches with the wt and Parp14 deficient mice specific gene detections using the Fischer’s Exact test and the FDR<0.05 filter (Fig. 4B, and Suppl. data file 1_7-12). When we looked at the GO Biological Process (BP) terms, identified based on the 325 genes specifically detected in the Parp14 deficient mice, all the 20 BP terms were related to cell division (Fig. 4C). Altogether 201 genes out of the 325 analyzed genes were mapped to these BPs (Suppl. data file 1_13). When we ran the GO term analysis with the 520 genes specifically detected in the wt mice, 23 BP terms were identified (Fig. 4C). Seven of these BP terms had relevance to infection and immune responses, e.g., neutrophil chemotaxis and leukocyte migration. Altogether 46 genes were behind these seven BP term identifications, e.g., cytokines *ccl17*, *ccl7*, *ccl2*, *cxcl9* and *cxcl10* (Suppl. data file 1_14). The *ccl2* gene was validated in a TaqMan qPCR analysis. We found out that expression of the *ccl2* gene was significantly higher in the wt mice, i.e., hampered in the Parp14 deficient mice (Fig. 5). Such an expression pattern was not detected in the small intestine or cecum (Fig S5). Next, we ran KEGG pathway analyses using the wt and Parp14 deficient mice specific gene detections. Based on the canonical <0.05 *p*-value filtering, only two KEGG pathways were detected with the Parp14 deficient mice specific gene list, and none were significant based on the <0.05 pAdj based filtering (Fig. 4D, Suppl. data file 1_16). In contrast, based on the canonical <0.05 *p*-value filtering, we identified 20 KEGG pathways with the wt specific genes (Fig. 4D, Suppl. data file 1_15) and one of these was significant based on the <0.05 pAdj based filtering. This pAdj significant KEGG pathway was the IL-17 pathway containing altogether 8 scored genes (*lcn2*, *il1b*, *s100a8*, *s100a9*, *cxcl10*, *ccl17*, *ccl7*, *ccl2*). The *il1b* gene was validated in a TaqMan qPCR analysis. We found out that expression of the *il1b* gene was significantly higher in the wt mice, i.e., hampered in the Parp14 deficient mice (Fig. 5). Such an expression pattern was not detected in the small intestine or cecum (Fig S5). Taken together, we detected a cell division related transcriptional signature in the Parp14 deficient mice that was missing from the wt mice in *S.* Typhimurium infection. Also, we detected an infection and immune response related transcriptional signature in the wt mice that was missing from the Parp14 deficient mice in *S.* Typhimurium infection.

**Figure 4.**
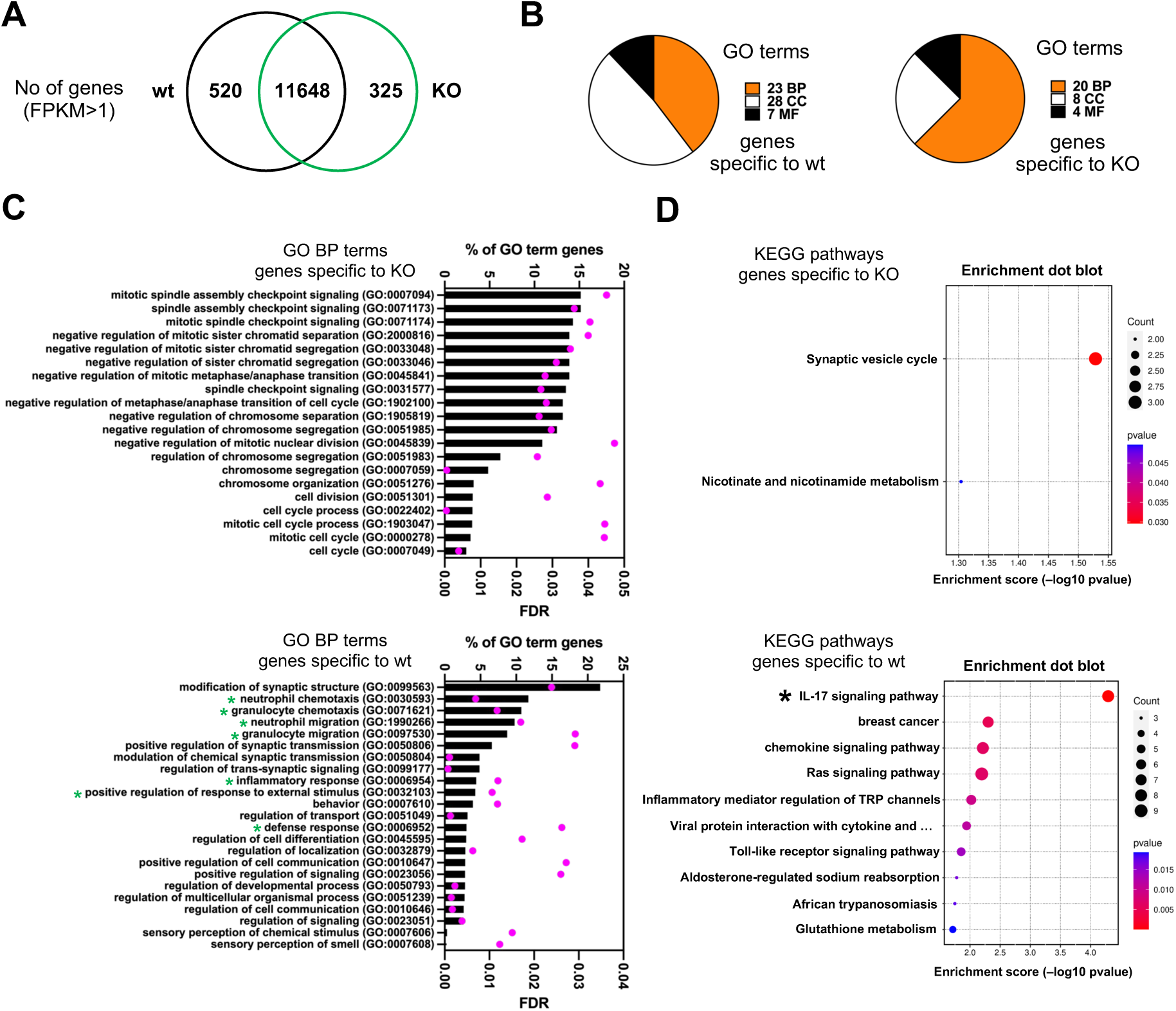
Transcriptional signatures uniquely detected in *S*. Typhimurium infected wt and Parp14 deficient mice. Data of a triplicate RNA-Seq analysis of mouse large intestine sections 1 day post-infection are shown. **A)** The Venn diagrams of shared and unique genes that were detected to be expressed in the infected wt and Parp14-deficient mice [Fragments Per Kilobase of transcript per Million mapped reads (FPKM)-value >1]. The integer is the number of genes detected to be expressed in both of the genotypes. **B)** The pie charts of the numbers of identified Gene Ontology (GO) terms based on the genotype specific lists of expressed genes (BP, biological process; CC, cellular component; MF, molecular function; Suppl. Data file 1). **C)** Bar graph representation of all the identified GO BP terms with the genotype specific lists of expressed genes. The BP terms are sorted based on the % of GO term genes-values (number of detected genes in a particular BP term / number of all genes in particular BP term x 100). FDR refers to the false discovery rate-value. A false discovery rate (FDR)-value cut-off of <0.05 was used in the searches. The green asterisks in the wt sub-panel refer to the seven infection and immune response related BP terms. **D)** Pathway enrichment dot blot representations of all (KO sub-panel) and the top 10 (wt sub-panel) Kyoto Encyclopedia of Genes and Genomes (KEGG) pathways identified with the genotype specific lists of expressed genes. All the identified KEGG pathways with the corresponding gene lists are described in Suppl. data file 1_15-16. The KEGG pathways are sorted based on the *p*-value. The count values refer to the number of genes that were detected in a particular KEGG pathway. The black asterisk in wt sub-panel refers to the KEGG pathway with a <0.05 pAdj-value.

**Figure 5.**
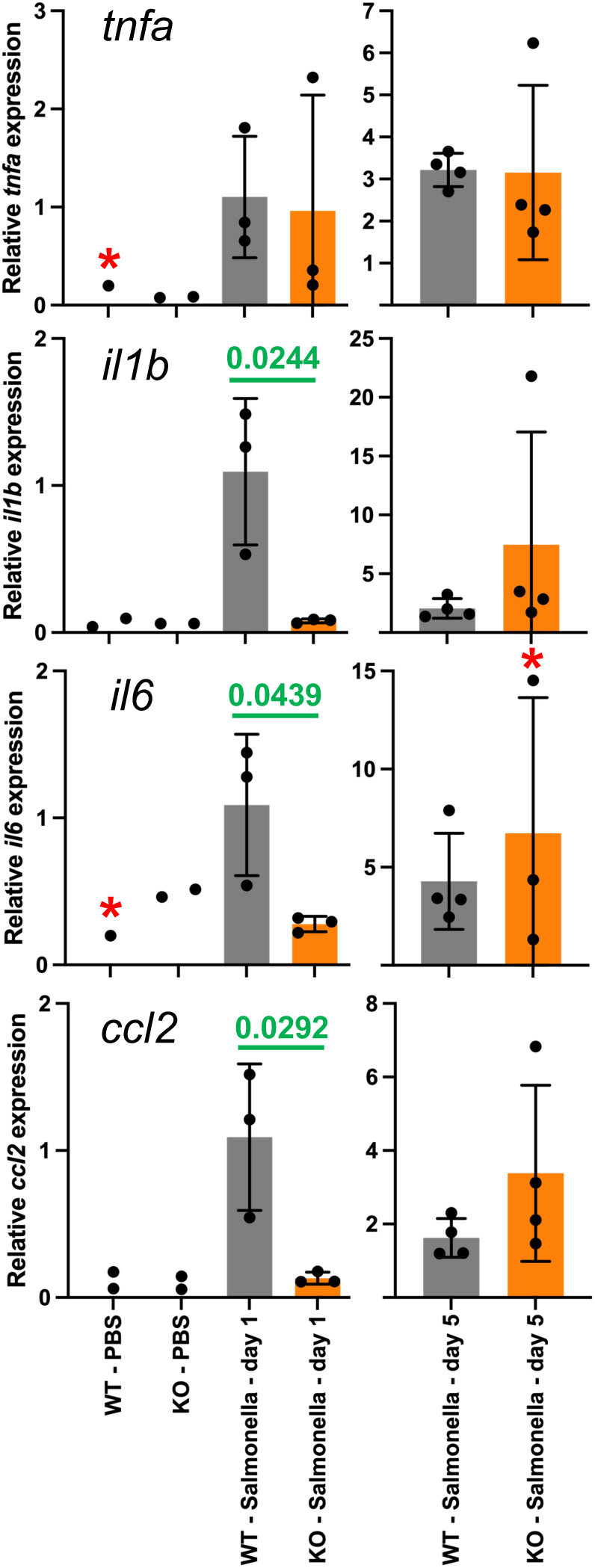
Hampered expression of three cytokines in the large intestine of *S.* Typhimurium infected Parp14 deficient mice. The panels display the TaqMan qPCR data on relative cytokine expression with means and standard deviation. Statistical analyses were done with two-tailed unpaired t-test to compare experimental conditions with 3 or more biological replicates. All the statistically significant differences between the wt and Parp14 deficient mice are indicated in the sub-panels. Samples were included in the displayed data analysis if they passed the 0.5 standard deviation Ct filter of replicate TaqMan runs. The red asterisks refer to experimental conditions where one or more samples did not pass this 0.5 standard deviation Ct filter. The calibrators in all sub-panels are the mean dCq-values of the day 1 infected wt mice.

### Transcriptional signature down-regulated in large intestine of *S*. Typhimurium infected Parp14 deficient mice

Next, we turned our attention to the detected differentially expressed genes (DEGs) in triplicate groups of day 1 infected Parp14 deficient vs. wt mice (Suppl. data file 1_17). The pairwise R2 Pearson correlation values of the FPKM-values of the DEGs were 0.974 or higher with the Parp14 deficient group, and 0.964 or higher with the wt group (Fig. 6A). This indicates that the triplicate samples could be reliably pooled for further analysis. The volcano plot of all the DEGs is shown in Fig. 6B. To execute downstream analysis with these DEGs, we ran the DEGs through pAdj<0.05 filter, which resulted into a radically shorter list of genes, i.e., 52 up-regulated genes, and 158 down-regulated genes (Suppl. data file 1_17-18). When we ran a GO analysis with the up-regulated genes, i.e., the genes that had more transcripts in the knockout as compared with the wt mice, using the canonical Fischer’s test and FDR<0.05 filter, we did not detect a single BP, CC or MF GO terms, Fig. 6C). In contrast, with down-regulated genes, i.e., the genes that had less transcripts in the knockout as compared with the wt mice, numerous BP, CC and MF GO terms were detected (Suppl. data file 1_19-21, Fig. 6C-D). Many of the top down-regulated DEG BP GO terms had functional relevance with cell adhesion and cytoskeleton remodeling (Fig. 6D). To further assess this finding, we performed a KEGG pathway analysis with the down-regulated DEGs. Based on the canonical <0.05 p-value filtering, we identified 12 KEGG pathways, and two of these were significant based on the <0.05 pAdj based filtering (Fig. 6E, Suppl. data file 1_22). The top pAdj significant KEGG hit was the Regulation of actin cytoskeleton-pathway containing altogether 8 scored genes (*ppp1r12b*, *pfn2*, *enah*, *myh11*, *fn1*, *rdx*, *myl9*, *myh10*). Many of these genes were shared with the lower ranking KEGG hits, including the Focal adhesion-, Tight junction- and *Salmonella* infection-pathways (Suppl. data file 1_22). Taken together, we detected a down-regulated cell adhesion and cytoskeleton remodeling related transcriptional signature in the *S.* Typhimurium infected Parp14 deficient mice.

**Figure 6.**
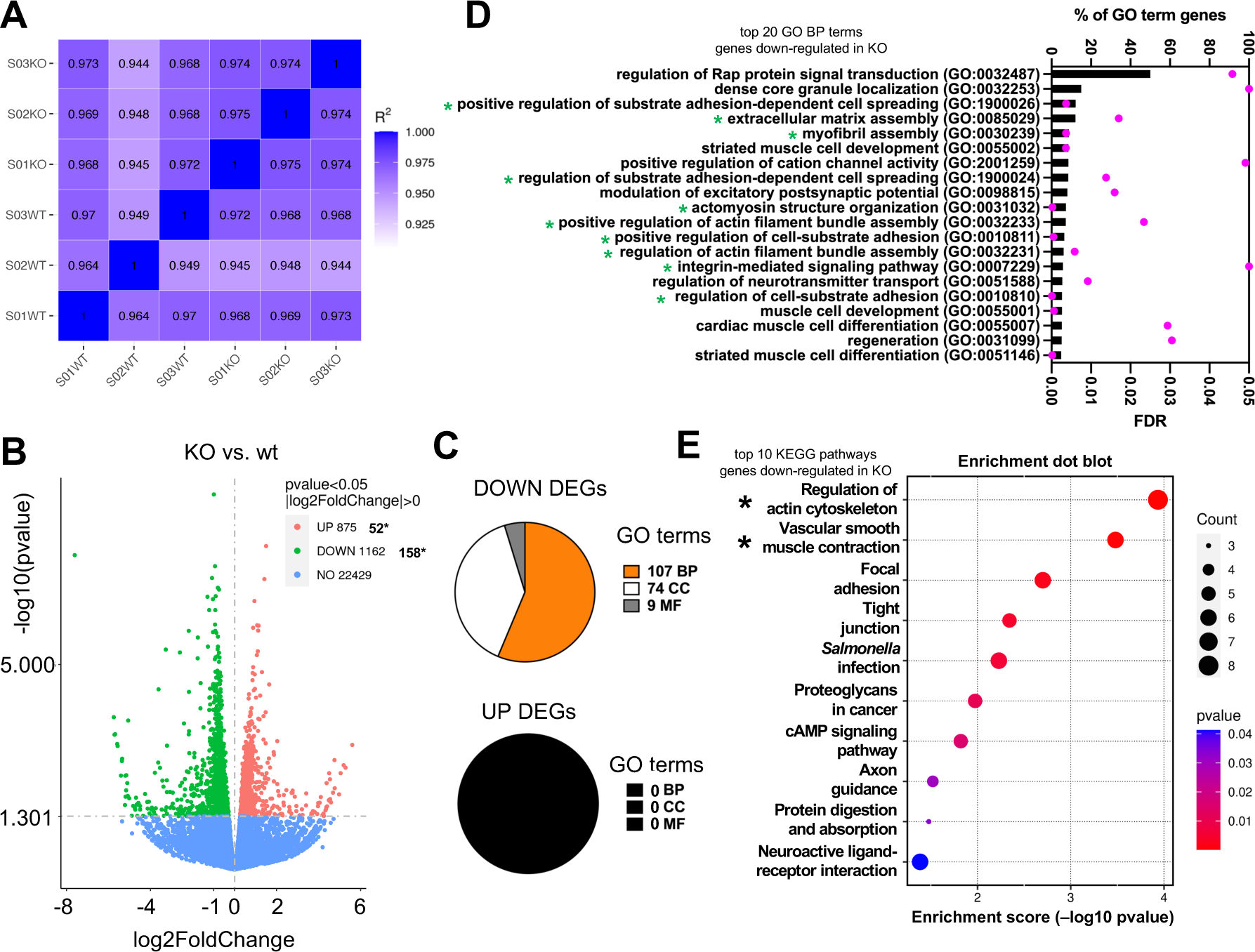
Transcriptional signature down-regulated in *S*. Typhimurium infected Parp14 deficient mice. Data of a triplicate bulk tissue RNA-Seq analysis of mouse large intestine sections 1 day post-infection are shown. **A)** Inter-sample correlation heatmap based on the Fragments Per Kilobase of transcript per Million mapped reads (FPKM)-values of the differentially expressed genes (DEGs) in Parp14 deficient vs. wt mice comparison. R^2^ is the square of Pearson correlation coefficient (R). **B)** Volcano plots of the DEGs. Specific information of the DEGs is given in Suppl. data file 1_17-18. The x-axis shows the fold difference in gene expression between different samples, and the y-axis shows the statistical significance of the differences. Red dots represent up-regulation genes and green dots represent down-regulation genes. The dashed line indicates the threshold line for statistically significant differential gene expression. The values marked with asterisks refer to the number of DEGs that were used for all the subsequent data analysis after filtering, i.e., UP genes, log2(FoldChange) > 0.5 & pAdj < 0.05; DOWN genes, log2(FoldChange) < −0.5 & pAdj < 0.05 (Suppl. data file 1_18). **C)** Gene Ontology (GO) term analysis with DEGs in Parp14 deficient vs. wt mice comparison (BP, biological process; CC, cellular component; MF, molecular function; Suppl. Data file 1_19-21). The GO terms were searched using the canonical Fischer’s test and a false discovery rate (FDR)-value <0.05 filter. **D)** Bar graph representations of the top 20 identified GO BP terms (all the 107 identified GO BP terms in Suppl. Data file 1-19) sorted based on the % of GO term genes-values (number of detected genes in a particular GO term / number of all genes in a particular GO term x 100). **E)** Pathway enrichment dot blot representations of the top 10 identified Kyoto Encyclopedia of Genes and Genomes (KEGG) pathways sorted based on the *p*-value. All the identified KEGG pathways with the corresponding gene lists are described in Suppl. data file 1-22. The count values refer to the number of genes that were detected in a particular KEGG pathway. The black asterisk in the wt sub-panel refers to the KEGG pathway with a <0.05 pAdj value.

## DISCUSSION

The microbiota-mediated colonization resistance of incoming pathogens in the gastrointestinal tract is supported by thick intestinal mucous layer secreted by the epithelial cells (9, 10). If these barriers are breached, a strong innate inflammatory response is initiated, followed by the adaptive immune system activation, all designed to eradicate the invading pathogen. We previously reported that the lack of Parp14 sensitized mice to dextran sulfate sodium (DSS) induced colitis, manifested, for example, as stronger epithelial erosion compared to the wt mice (35). Oral administration of the non-physiological DSS chemical to mice via drinking water, used frequently to model human ulcerative colitis (46, 47), induces severe colitis characterized by weight loss, bloody diarrhea, loss of epithelial cells and infiltration of neutrophils. DSS is believed to cause direct damage to epithelial cells followed by strong innate immune system activation by dissemination of the proinflammatory colon luminal content such as bacteria into the sub-epithelial space (46, 47). Most likely, the strong pro-inflammatory response including oxidative stress exacerbates the deleterious effects of DSS. Here, we set out to study the expression and function of Parp14 in a bacterial enteropathogen mouse model.

We used the mouse streptomycin-pretreatment model of *S.* Typhimurium colitis (8, 33). The streptomycin treatment transiently depletes the microbiota and the subsequent *S.* Typhimurium colitis resembles many aspects of human disease, for example, epithelial erosion and massive infiltration of neutrophils (33). First, we analyzed the expression pattern of Parp14 upon infection. Histological examination demonstrated that the lamina propria of small intestine, cecum and large intestine contained Parp14 expressing cells. Based on the staining for the classical F4/80 macrophage marker, some of the Parp14-positive cells were macrophages. This resembles what we and others previously reported for the large intestine in the oral DSS exposure mouse model of IBD, and in IBD patients (35, 48). In the large intestine it appeared that the most strongly Parp14-staining F4/80-positive cells were frequently embedded in the epithelial cell layer having protrusions towards the gut lumen. Contacts with the luminal content might have driven the detected strong expression of Parp14, e.g., with bacterial lipopolysaccharide (LPS), which is a known inducer of Parp14 expression in human and mouse macrophages *in vitro* (32, 35). Overall, the strongest Parp14 staining was witnessed in the epithelial cells. To substantiate the epithelial cell findings, we analyzed the previously published single cell RNA-Seq data of eight different small intestine epithelial cell populations in the control and *S.* Typhimurium infected mice (37). We found out that the expression of *parp14* in the infected mice was pronounced in the enterocytes and Tuft cells. The latter ones are rare and functionally elusive cells found in the mucosal epithelia with immunomodulatory potential (49). It appears possible that Parp14 has specific infection-associated functions in the Tuft cells, in addition to the dominant intestinal epithelial cell population, that is, the enterocytes. We also quantified Parp14 immunohistochemical staining in the epithelial cells with QuPath throughout the gastrointestinal tract and detected tissue specific temporal patterns. One of the most interesting patterns was detected in the large intestine. *Salmonella* infection increased the nuclear Parp14 staining at day 1, but, at day 5, less staining was detected, although the overall cellular Parp14 staining increased from the resting state to day 1 and further to day 5. We interpret this as a sign of an early infection requirement for nuclear Parp14. Most likely, this finding reflects the potent role of Parp14 as a transcriptional regulator, which has been studied so far in lymphocytes and macrophages (17, 18, 27, 32). For example, Parp14 accumulates to the nucleus of LPS-stimulated RAW264.7 macrophages *in vitro*, and influences the transcriptional response to LPS stimulation (32). Overall, Parp14 appears to be expressed throughout the gastrointestinal tract in *S.* Typhimurium infection in macrophages, but, in particular, in the epithelial cells.

We detected significant effects of Parp14 deficiency on the severity of salmonellosis. Out of the measured variables at day 1 and day 5 post-infection, the mouse weight, colon length, spleen weight and liver weight were similar in the infected wt and Parp14 deficient mice. Also, the numbers of viable bacteria were mostly similar between the mouse genotypes in small intestine, large intestine, mesenteric lymph nodes, spleen and liver. However, we witnessed increased numbers of viable bacteria in the liver of Parp14 deficient mice at day 1. At the same time point, we saw that the small intestine of Parp14 deficient mice had stronger epithelial erosion. We are tempted to speculate that the poor condition of the small intestine epithelium had contributed to the enhanced peripheral tissue invasion of *S.* typhimurium. In part, this effect could include defective Peyer’s patches and M cells therein, which are well-known entry sites for peripheral tissue invasion of *S.* Typhimurium (50, 51). Interestingly, stronger epithelial cell erosion was not witnessed in the infected Parp14 deficient mice at day 5. Therefore, the deleterious effect of Parp14 deficiency in the small intestine appeared to be transient. In part, our QuPath-based detection of higher Parp14 staining in the small intestine epithelial cells at day 1 as compared to the resting state or day 5 support the notion of functional importance for Parp14 in the early timepoints of infection. At day 5, we witnessed lower amounts of viable bacteria in the large intestine of Parp14 deficient mice. At the same time, we saw that the large intestine of Parp14 deficient mice had stronger epithelial erosion, immune cell infiltration and Goblet cell loss. These day 5 findings could imply that the mucosal inflammation in the infected Parp14 deficient mice was hyperactive resolving the infection more efficiently, but, at the same time, it drove tissue destruction in the large intestine. Overall, it appears that Parp14 is involved in the regulation of mucosal inflammation in *S.* Typhimurium infection or it influences the mucosal epithelial barrier integrity, or possibly both of these phenomena.

To possibly obtain mechanistic insights on the exacerbated salmonellosis in Parp14 deficient mice, we executed a bulk tissue RNA-Seq analysis of the large intestine. Based on the GO, but not on the KEGG analysis, we detected a cell cycle related transcriptional signature in the infected Parp14 deficient mice that was missing from the wt mice. We interpret this as a sign that the infected Parp14 deficient mice had problems in or an increased need for cellular renewal in the large intestine. We also detected an infection and immune response related transcriptional signature in the wt mice that was missing from the Parp14 deficient mice. Seven of the 23 identified GO BP terms were related to infection and immune responses with altogether 46 genes. Moreover, we identified one KEGG pathway, that is, the IL-17 signaling pathway, which is a key regulatory pathway of mucosal inflammation (52). There were 8 genes behind this KEGG hit (*lcn2*, *il1b*, *s100a8*, *s100a9*, *cxcl10*, *ccl17*, *ccl2*, and *ccl7*), all of them also scoring the seven infection and immune response related GO BP terms. Importantly, when the gene lists of the seven infection and immune response related GO BP terms were compared with the duplicate PBS wt control mice gene lists, we found that only 4 genes were shared (*hp*, *gprc5b*, *trim30a* and *h2-Q10*). This indicates that the unique detection of most of the 46 genes in the infected wt mice was indeed associated with the infection, not merely with the mouse genotype difference. It is tempting to speculate that the hampered downstream effector functions of the IL-17 signaling pathway contributed to the exacerbated salmonellosis in Parp14 deficient mice. For example, it is known that a more severe *S.* Typhimurium infection develops in mice deficient of the chemokine Ccl2, encoded by the *ccl2* gene (53). Also, the metal chelating calprotectin heterodimer, encoded by the *s100a8* and *s100a9* genes, is known to directly inhibit the growth of *S.* Typhimurium *in vitro* (54). Overall, it appears, based on the RNA-Seq functional proximations, that the *S.* Typhimurium infected Parp14 deficient mice had problems to mount a proper innate immune response in the large intestine.

Interesting down-regulated transcriptional signatures were detected in the *S.* Typhimurium infected Parp14 deficient mice. Based on the analysis of 158 downregulated genes, several GO BP terms having functional relevance with cell adhesion and cytoskeleton remodeling were identified. Moreover, two KEGG pathways were identified, that is, the regulation of actin cytoskeleton pathway and the vascular smooth muscle contraction pathway. Many of the genes behind these two KEGG hits, e.g., *pfn2*, *fn1*, *rdx*, *myh10-11* and *myl9*, were also shared with the lower ranking >0.05 pAdj KEGG pathways, including the focal adhesion pathway and the tight junction pathway. These RNA-Seq findings potentially reflect the earlier *in vitro* indications of hampered cell motility and cell adhesion paralleled with abnormal F-actin staining in the absence of Parp14 (55, 56). It has remained unknown how Parp14 could mechanistically regulate cell motility and cell adhesion, although localization of Parp14 to the focal adhesion complexes has been detected (55, 57). Our data imply that Parp14 could also shape the abundancies of proteins functionally important for cell motility and cell adhesion via transcriptional regulation. However, more experimental work is required as it is also plausible that we were observing an adaptive cellular response to the lack of a more direct mechanical cytosolic function of Parp14, e.g., at the focal adhesion complexes. Overall, it appears, based on the RNA-Seq functional proximations, that *S.* Typhimurium infected Parp14 deficient mice had problems in the cell motility and cell adhesion processes in the large intestine.

In summary, our study with body-wide Parp14 deficient mice (34, 35) provides compelling *in vivo* evidence that Parp14 has functional relevance in the physiological response to a bacterial infection. To the best of our knowledge, only one previous *in vitro* phenotypic study employing Parp14 depletion in bacterial infection has been published (32). The Parp14 deficient RAW264.7 macrophages contained more viable *S*. Typhimurium bacteria, and, upon *S*. Typhimurium infection, produced less microbicidal nitric oxide as well as had a defective expression pattern of a number of inflammation-related genes, such as *ifnb1*, *ccl5*, *cxcl10*, and *ifit1* (32). Indeed, macrophages might have been malfunctional in our *S.* typhimurium infected Parp14 deficient mice. However, based on our current and previous work (35), it appears that Parp14 has probable functions also in the epithelial cells. Further animal experimentation would be needed to resolve these issues, e.g., with cell lineage specific Parp14 knockout mice in parallel with single cell RNA-Seq studies, in order to identify the exact phenotype contributing Parp14 expressing cell type(s) and their specific functions in bacterial infection.

## Notes

### Competing Interest Statement

The authors have declared no competing interest.

