## Supplementary figures_1-5 for "Exacerbated salmonellosis in poly(ADP-ribose) polymerase 14 deficient mice"

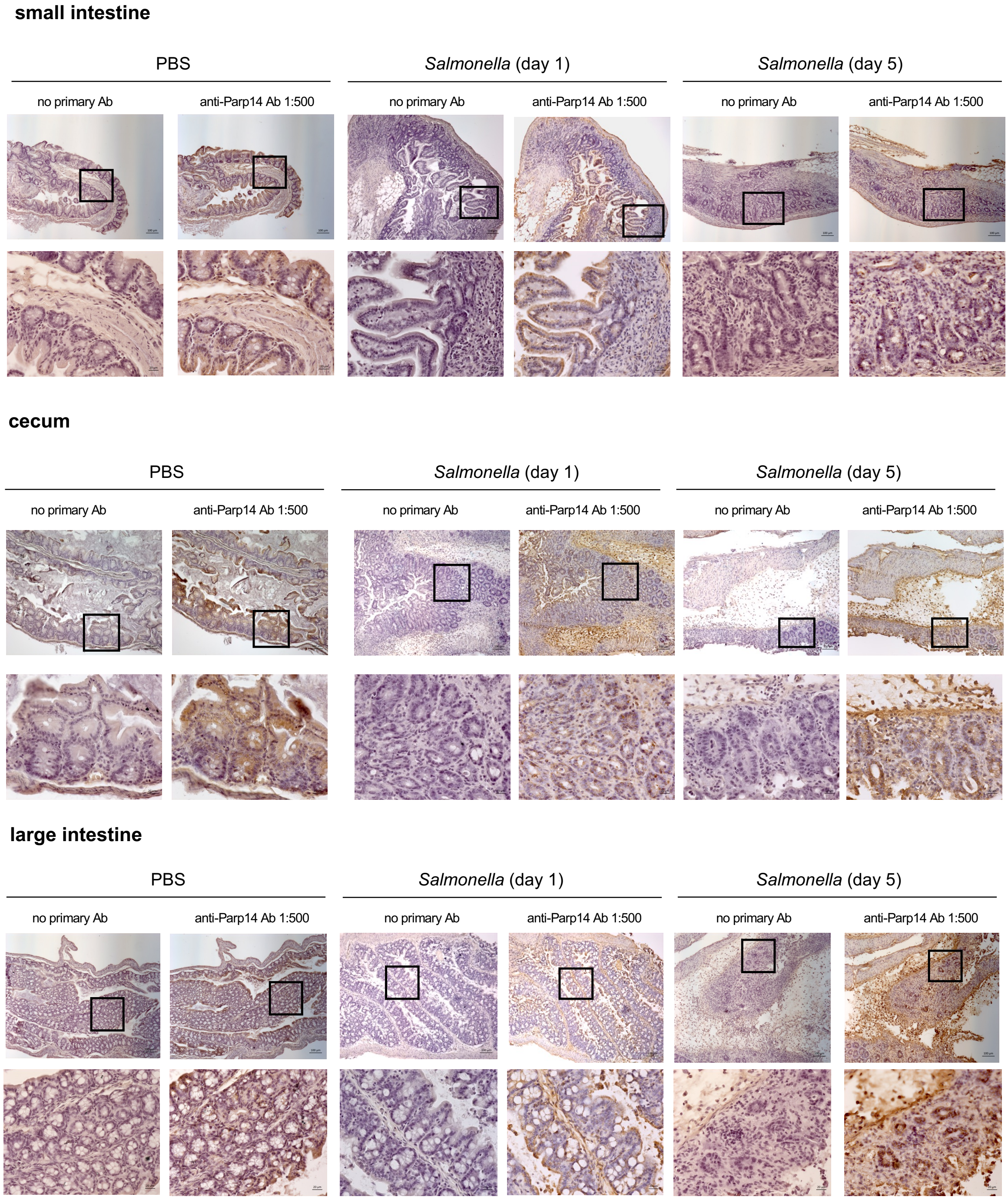


**Figure S1. Vedantham et al.**

**Figure S1. Immunohistochemical staining of Parp14 in the mouse gastrointestinal tract FFPE tissue sections.** For each gastrointestinal tract location, 10x air (scale bar, 100 μm) and 40x oil (scale bar, 20 μm) objective images are shown (1:500 dilution of the anti-Parp14 antibody, sc-377150, Santa Cruz Biotechnology).

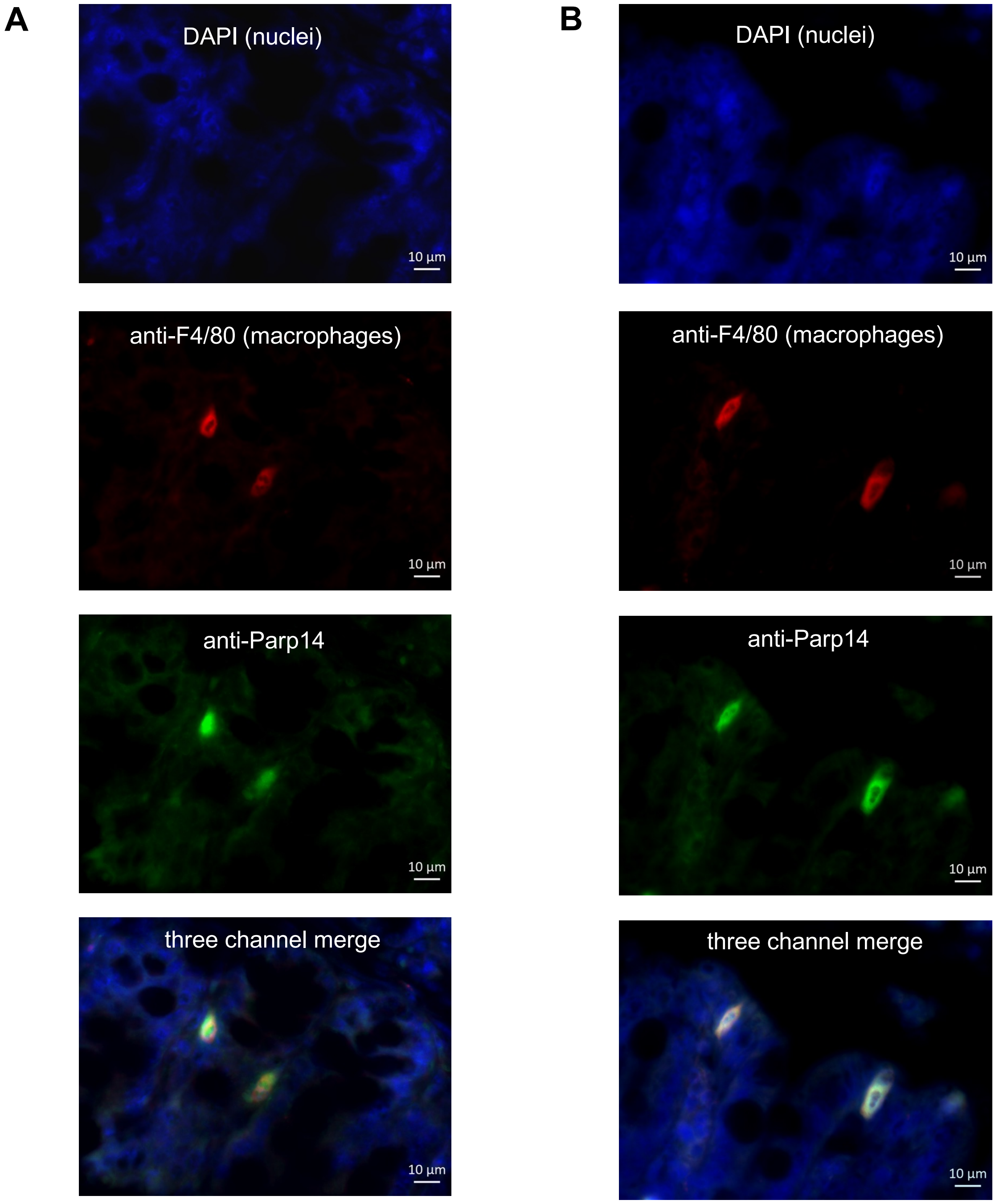


**Figure S2. Vedantham et al.**

**Figure S2. Double immunofluorescence staining of Parp14 and the macrophage marker F4/80 in the mouse large intestine FFPE tissue sections.** Two different tissue sections derived from the *Salmonella* infected C57BL/6N mice (day 1) were analyzed for Parp14 (green channel, 1:500 dilution of anti-Parp14 antibody) and macrophage marker F4/80 (red channel). **A)** Apparent location of Parp14 positive macrophages in the lamina propria. 63x objective image is shown (scale bar, 10 μm). **B)** Apparent location of Parp14 positive macrophages in the epithelial cell layer. 63x objective image is shown (scale bar, 10 μm).

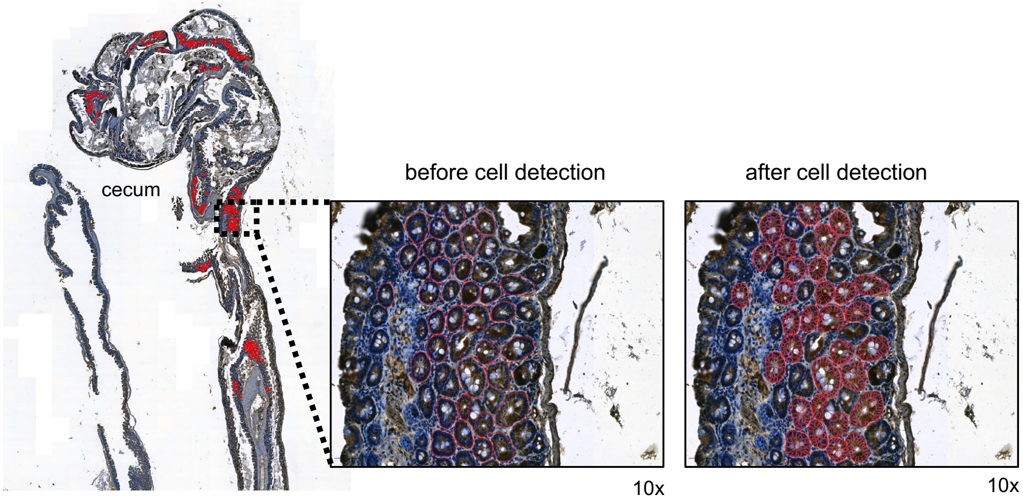


**Figure S3. Vedantham et al.**

**Figure S3. QuPath-based quantitation of Parp14 staining in the mouse gastrointestinal tract FFPE tissue sections.** Parp14 staining intensity was quantified with entire tissue sections by selecting 50-200 horizontal villus cross-sections (circled in red, before cell detection) and thereby thousands of individual epithelial cells (circled in red, after cell detection) per animal. The figure is an example of one cecum section quantitation.

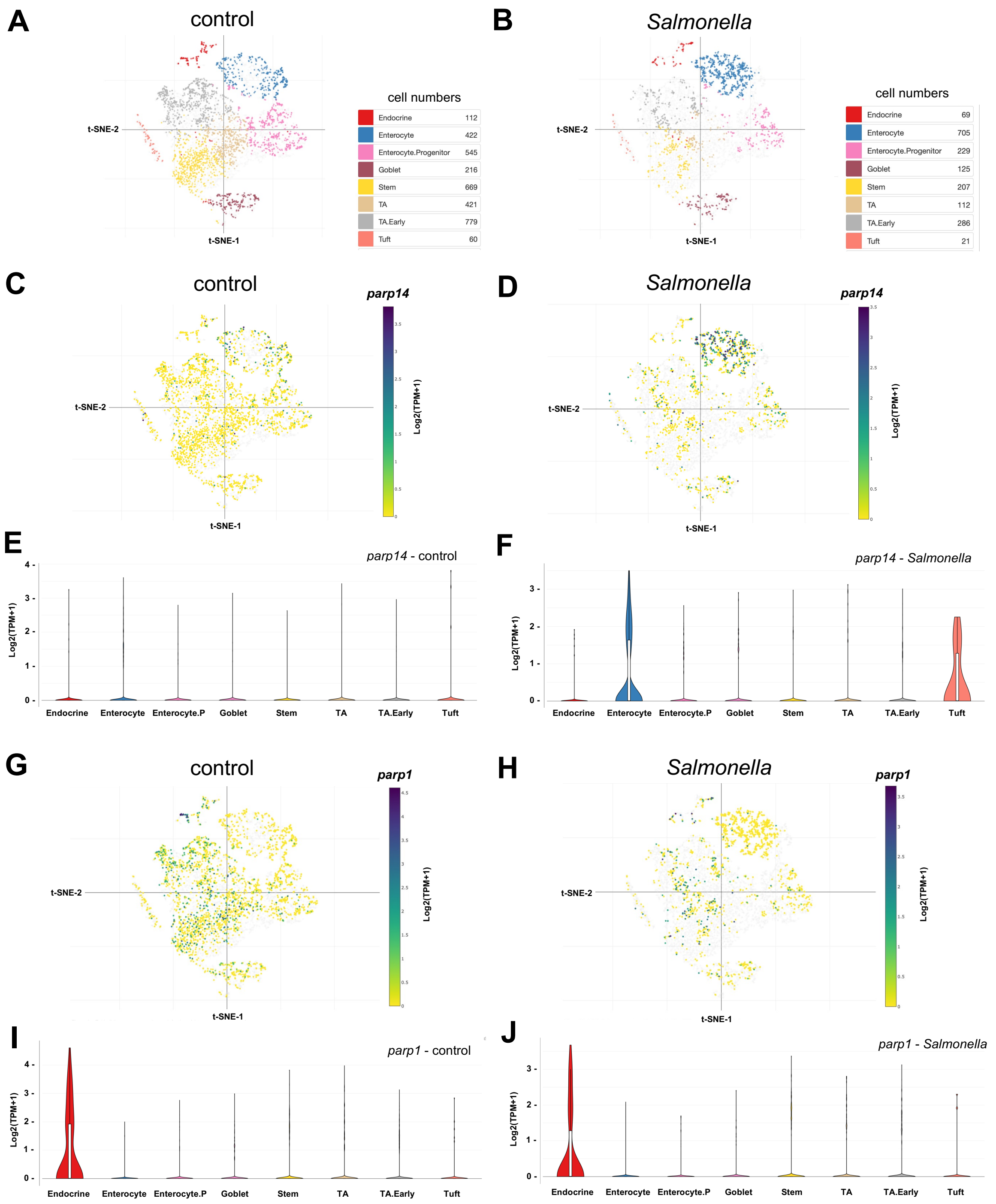


**Figure S4. Vedantham et al.**

**Figure S4. Single cell RNA-Seq analysis of *parp14* expression in different mouse epithelial cell subtypes of the small intestine. A-B)** Description of the identified mouse epithelial cell subtypes and their numbers in control vs. *Salmonella*-infected mice 2 days post-infection. The t-distributed stochastic neighbor embedding (t-SNE) method was used to visualize the data. **C-D)** Expression levels of *parp14* in the different epithelial cell subtypes. The expression of *parp14* is displayed in Log2(TPM+1)-values, that is, log2-transformed transcript per million-values. The t-distributed stochastic neighbor embedding (t-SNE) method was used to visualize the data. **E-F)** Distribution blot of *parp14* expression levels in the different epithelial cell subtypes. **G-H)** Expression levels of *parp1* in the different epithelial cell subtypes. The expression of *parp1* is displayed in Log2(TPM+1)-values, that is, log2-transformed transcript per million-values. The t-distributed stochastic neighbor embedding (t-SNE) method was used to visualize the data. **I-J)** Distribution blot of *parp1* expression levels in the different epithelial cell subtypes. All the data was analyzed and visualized using the single cell RNA-Seq data analysis and visualization interface at the Broad Institute Single Cell Portal (https://singlecell.broadinstitute.org/single_cell).

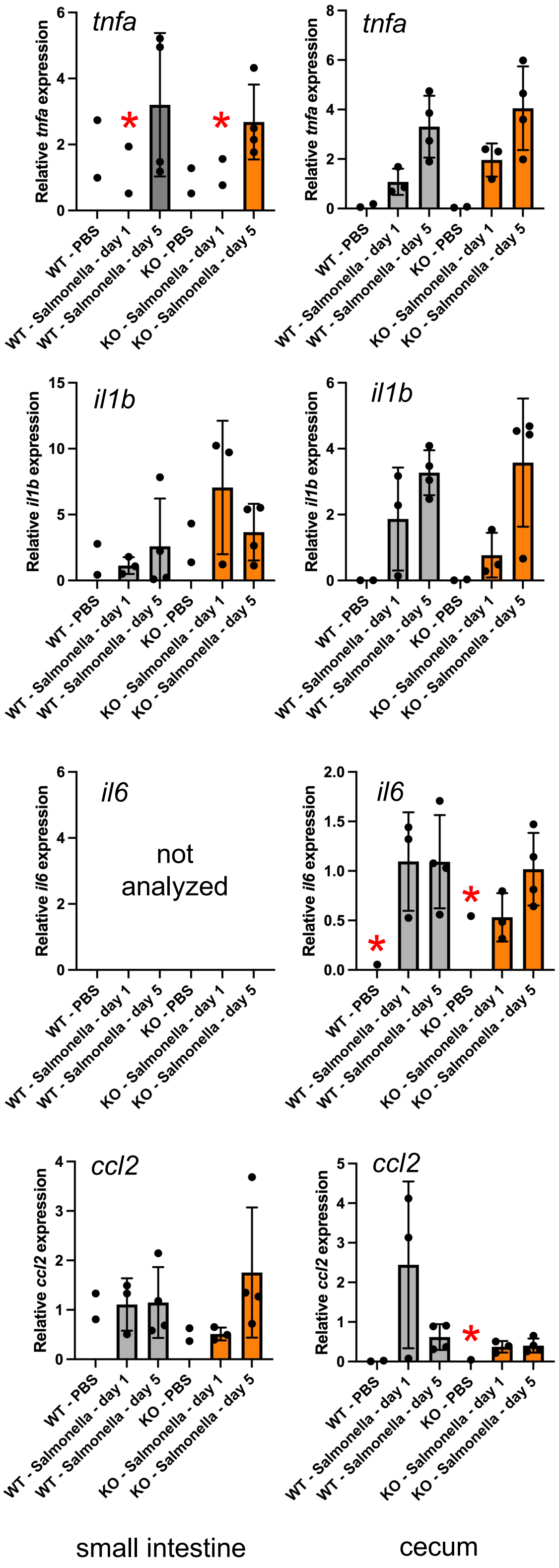


**Figure S5. Vedantham et al.**

**Figure S5. TaqMan qPCR-based quantitation of cytokine expression in the cecum and small intestine.** The panels display the TaqMan qPCR data on relative cytokine expression with means and standard deviation. Statistical analyses were done with two-tailed unpaired t-test to compare experimental conditions with 3 or more biological replicates. No statistically significant differences between the wt and Parp14-deficient mice were detected. Samples were included in the displayed data analysis if they passed the 0.5 standard deviation Ct filter of replicate TaqMan runs. The red asterisks refer to experimental conditions where one or more samples did not pass this 0.5 Ct filter. The calibrators in all sub-panels are the mean dCq-values of the day 1 *Salmonella* infected wt mice.
